# Linking light-dependent life history traits with population dynamics for *Prochlorococcus* and cyanophage

**DOI:** 10.1101/696435

**Authors:** David Demory, Riyue Liu, Yue Chen, Fangxin Zhao, Ashley Coenen, Qinglu Zeng, Joshua S. Weitz

## Abstract

*Prochlorococcus* grow in diurnal rhythms driven by diel cycles. Their ecology depends on light, nutrients, and top-down mortality processes including lysis by viruses. Cyanophage, viruses that infect cyanobacteria, are also impacted by light. For example, extracellular viability and intra-cell infection kinetics of some cyanophage vary between light and dark conditions. Nonetheless, it remains unclear if light-dependent viral life history traits scale-up to influence population-level dynamics. Here we examined the impact of diel-forcing on both cellular- and population-scale dynamics in multiple *Prochlorococcus*-phage systems. To do so, we developed a light-driven population model including both cellular growth and viral infection dynamics. We then tested the model against measurements of experimental infection dynamics with diel forcing to examine the extent to which population level changes in both viral and host abundances could be explained by light-dependent life history traits. Model-data integration reveals that light-dependent adsorption can improve fits to population dynamics for some virus-host pairs. However, light-dependent variation alone does not fully explain realized host and virus population dynamics. Instead, we show evidence of a previously unrecognized lysis saturation at relatively high virus to cell ratios. Altogether, our study represents a quantitative approach to integrate mechanistic models to reconcile *Prochlorococcus*-virus dynamics spanning cellular to population scales.

## Introduction

The unicellular cyanobacterium *Prochlorococcus* dominates the phytoplankton community and is a major contributor to primary production in tropical and subtropical oligotrophic oceans [1]. The ecology of *Prochlorococcus* is a function of physicochemical properties of the marine environment [2–4], bottom-up (i.e., nutrient driven) as well as positive top-down (i.e., mortality-driven) processes [5–11]. Amongst top-down factors, cyanophage (i.e., viruses that infect cyanobacteria) are highly abundant, and can be responsible for up to 30% of mortality in marine environments [12–17]. Light, temperature and nutrients influence *Prochlorococcus* growth [3, 4, 18] as well as its interactions with cyanophage [19].

*Prochlorococcus* are distributed across temperature and light gradients in the ocean environment [3, 20–22]. They are specialized into High-Light (HL), and Low-Light (LL) adapted ecotypes [23–25]. LL ecotypes have a high fluorescence and photo-inhibited growth at medium light intensity. They grow faster at low irradiance with a high concentration of divinyl chlorophyll *a* and *b* and have several *pcb* genes encoding constitutive photosystems I and II [23–25]. In contrast, HL ecotypes grow faster at medium light intensities, have a low concentration of divinyl chlorophyll *a* and *b*, and have only constitutive photosystem II light-harvesting complexes [23–25]. *Prochlorococcus* cells do not have a circadian rhythm but rather a diurnal rhythm that persists and can be synchronized under light-dark cycles [22, 26, 27]. This diurnal rhythm is divided into photosynthesis during the light phase and cell division associated with energy consumption during the dark phase [21, 28].

Cyanophage are also impacted by light. Ultraviolet radiation (UV) can lead to viral inactivation and degradation in the environment [19] by degrading proteins and altering viral structure [29, 30]. However, light can also affect viral interactions with and inside host cells. Some studies suggested a possible dependence of viral production on the host cell cycle in different phytoplankton lineages [31–34] whereas the intracellular production of some viruses may be decoupled from host cell cycle and light levels [35, 36]. A recent paper on the diel infection pattern of cyanophage infecting *Prochlorococcus* [37] suggests that the adsorption rhythm, as well as the transcription rhythm of cyanophage, may be related to the light-dark cycle and not to the host cell cycle. Analyzing light impacts on cyanophage-cyanobacteria dynamics requires some elaboration of the cyanophage life cycle.

The lytic cyanophage cycle can be summarized into three phases: the adsorption phase where virions attach to their host and inject their genetic material into the host cell, the cyanophage replication phase using the host machinery and the lytic phase where new viruses (virions) are released by lysing their host (see Figure 1a). Light can affect each of these phases and associated viral life history traits (LHTs). A study on *Synechococcus* infection showed a significant decrease in adsorption in the dark condition for some phages [36, 38, 39], whereas other cyanophage adsorb during light or dark conditions [36]. Similarly, light conditions also modify the virus cycle during the cyanophage replication phase. A positive relationship between light and viral production and reduction at dark have been reported for *Synechococcus* [39–41] and *Prochlorococcus* [42–44]. Some cyanophage also have genes that encode proteins involved in photosynthesis, suggesting that they can maintain the photosynthetic activity of their host during infection [45, 46].

**FIG. 1:**
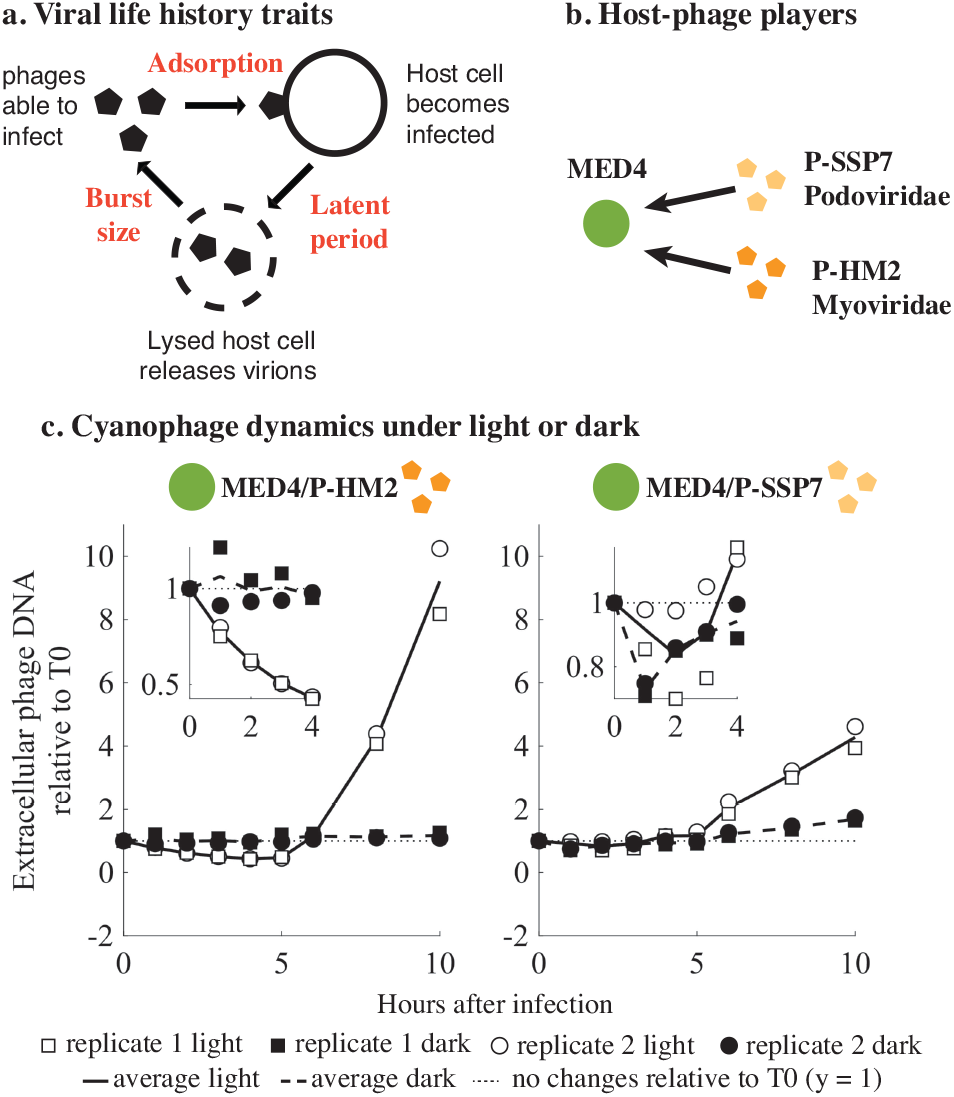
Cyanophage infection in the light or the dark. (a) Viral life history traits definition: viral adsorption (encounter and adsorption on a non-infected host, in mL hours-1)), latent period (time between adsorption and lysis of the host cell, in hours) and burst size (new phages produced per one lysed host cell). (b) Host-cyanophage pairs used in the study. (c) Infection under light or dark (adapted from [37], see methods). Cyanophage P-HM2 and P-SSP7 were used to infect their host cells under continuous light (empty symbol and solid line) or in the dark (filled symbol and dashed line). For all the host-phage pairs, the phage/host ratio is 0.1. Extracellular phage concentrations were measured as phage DNA by quantitative PCR and normalized to that at time 0 hour.

A salient example is that of Liu *et al.* [37] who investigated infection dynamics for cyanophage infecting *Prochlorococcus* under light-dark cycles (Figure 1b and c). The results suggest that cyanophage strains respond differently to light or dark conditions (Figure 1c). Infection under light was always efficient for all strains. However, P-SSP7 could infect and produce viruses in the dark while P-HM2 could not due to a lack of adsorption. These observations in fixed light or dark conditions form the central motivation for our study. That is: do differences in the response of viruses to light conditions at the cellular level explain population level dynamics of both *Prochlorococcus* and cyanophage given diurnal rhythms of light-dark cycles?

Here, we couple mathematical models, high-resolution (i.e., sub-daily) measurements, and model-data integration to explore the interactions between *Prochlorococcus* MED4 (a HL ecotype) with the cyanophage P-HM2 and P-SSP7 (Figure 1b). The models extend the framework of nonlinear population dynamics of lytic viruses and their hosts [47] to an explicitly light-driven context (see the related work of [48] on Cocolithoviruses and their *Emiliana huxleyi* hosts given low-resolution (i.e., daily) measurements). Our results suggest that cyanophage have light-dependent and host growth-dependent infection patterns with different associated life history traits depending on the viral strain. As we show, although diel-driven viral life history traits help explain population dynamics, they are not necessarily sufficient. Instead, our study identifies additional mechanisms involving saturating lysis that help reconcile population-level dynamics of cyanophage and cyanobacteria.

## Results

### Light-driven *Prochlorococcus* growth

We first estimated the growth of *Prochlorococcus* strain MED4 in culture under light-dark cycles and nutrient non-limited conditions during the exponential growth phase. We used an ordinary differential equation (ODE) model to describe the dynamics of the *Prochlorococcus* population (cells/ml):

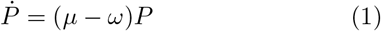

where *ω* is the host mortality rate (in h^−1^) and *μ* is the host growth rate (in h^−1^), as a function of perceived light during the experiments [49]:

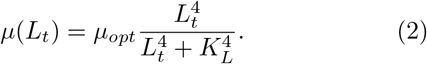

Here, *K_L_* is the minimum amount of light necessary to divide (in *μmol Quanta*) and *L_t_* is the cumulative light perceived by a cell at time *t* (in *μmol Quanta*) depending on the light-dark cycle state:

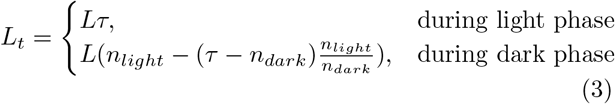

where, *τ* is equal to the remainder of the division of the time *t* by 24 hours (formally, *τ* = rem(*t*, 24)), *n_light_* and *n_dark_* are the number of hours of light and dark during the cycle respectively and *L* is the irradiance during the light phase of the experiments (in *μmol Quanta s*^−1^). *μ_opt_* is the optimal host growth (in *h^−1^)* rate defined by the growth-irradiance function described in [50] with the following equation (Eq. 4):

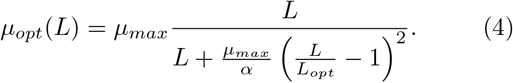

In this functional form, *μ_max_* is the maximal host growth rate (in *h*^−1^) at optimal light *L_opt_* (in *μmol* Quanta *s*^−1^) and *α* is the initial slope of the light response curve (in *h*^−1^). During the 24 hours of a light-dark cycle, *μ*(*L_t_*) increases during the first 14 hours (light period) from 0 to reach a maximum at *n_light_* and decrease during the *n_dark_* hours of the dark period.

The model in Eq. (1) was fit to population dynamic measurements of *Prochlorococcus* strain MED4 under a light-dark cycle [51] using a Markov Chain Monte Carlo approach (MCMC; see Methods). The best-fit light-driven host growth model recapitulates the experimental data (Figure 2a) with a good convergence of the MCMC parameter chains (Supplementary Figure 1) and Supplementary Table I). MED4 has a low growth-irradiance curve slope (*α* = 0.0011 h^−1^), a high optimal growth irradiance (*L_opt_* = 44.78 *μ* mol^−1^ Quanta^−1^ s^−1^) and a maximum growth rate of 0.0035 h^−1^ (Figure 2b and Supplementary Figure I. These parameters are consistent with prior estimates of HL growth-irradiance characteristics for strain MED4 [52] (Figure 2c).

**FIG. 2:**
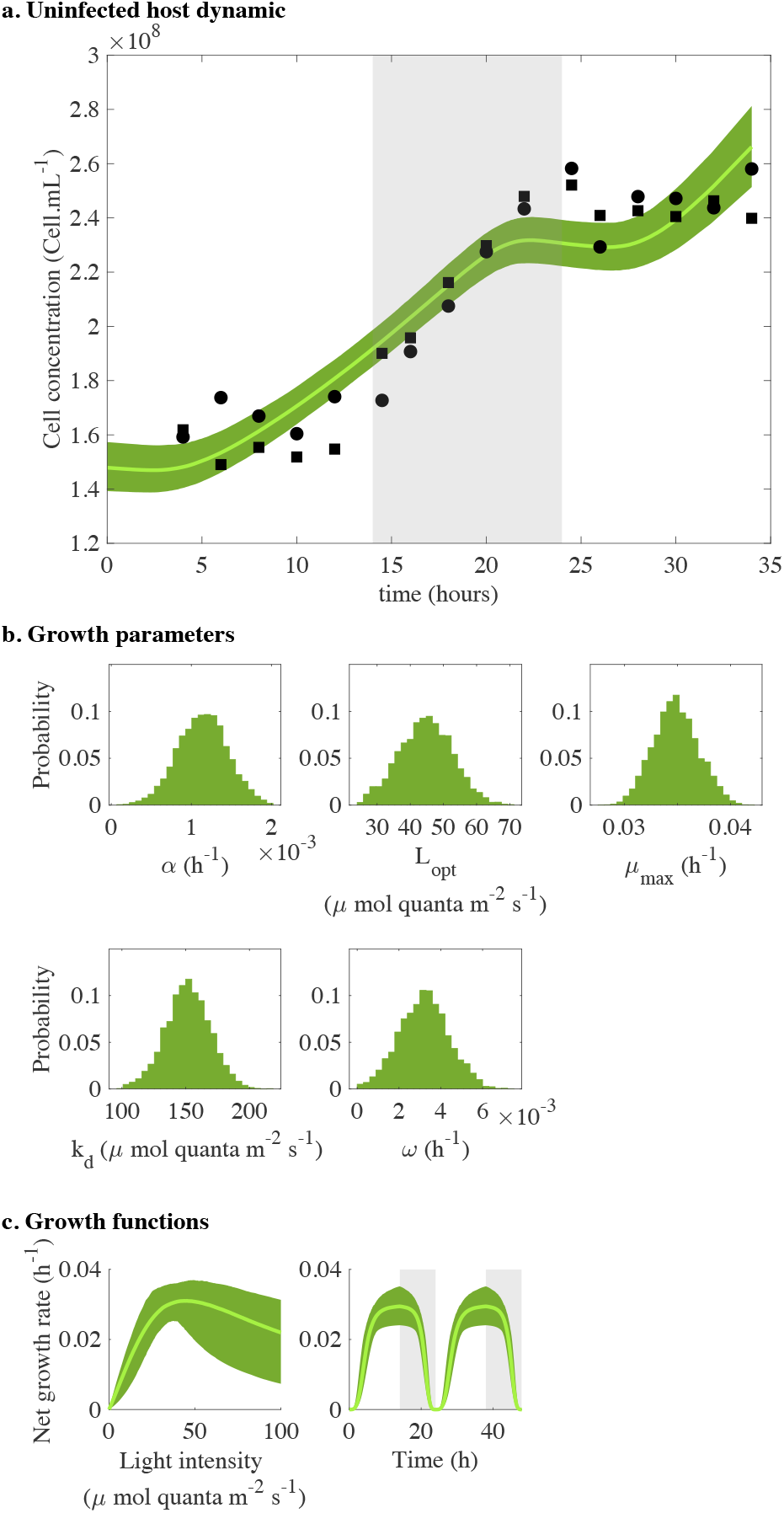
Modeling *Prochlorococcus* MED4 strain as function of light without viruses during the exponential phase. (a) Fit of the host dynamic (Eq. 1). Solid lines represent the median of 5000 model simulations and shade areas are the 95% quantiles. Black dots are data (from [51]) for two replicates and gray shaded area is the dark condition. (b) Growth parameters distributions of the host model (Eq. 1 and 4). Parameters distribution histograms estimate using a MCMC algorithm: PI-curve slope of the linear phase *α*, Optimal growth light *L_opt_*, Maximal growth rate *μ_max_*, Minimum amount of light necessary to divide *K_d_* and Natural mortality *ω*. (c) Growth functions that drive the host dynamic: growth is expressed as the net growth rate (*μ_opt_ − ω*) as function of irradiance (Eq. 4 - left panel) and as function of time (Eq. 2 - right panel).

### Modelling *Prochlorococcus*-phage dynamics under light-dark cycles

To investigate *Prochlorococcus*-cyanophage dynamics under light-dark cycles, we developed a nonlinear ODE population model describing the infection of *Prochlorococcus* by cyanophage (Figure 3a), extending existing frameworks for modeling obligately lytic phage-host interactions [47]. The host population is categorized as susceptible (S), exposed (E), and infected (I), such that the total host population is *N* = *S* + *E* + *I*. The density of free cyanophage is denoted by *V*. The model isdescribed by the following system:

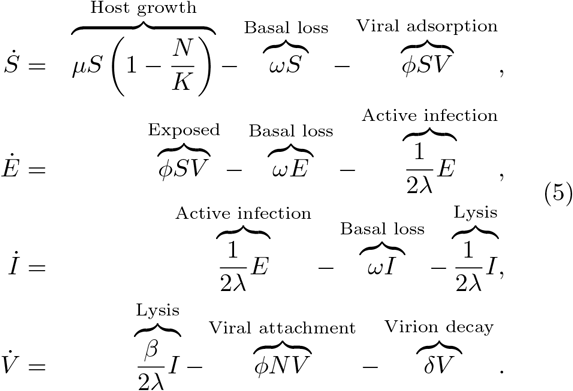

**FIG. 3:**
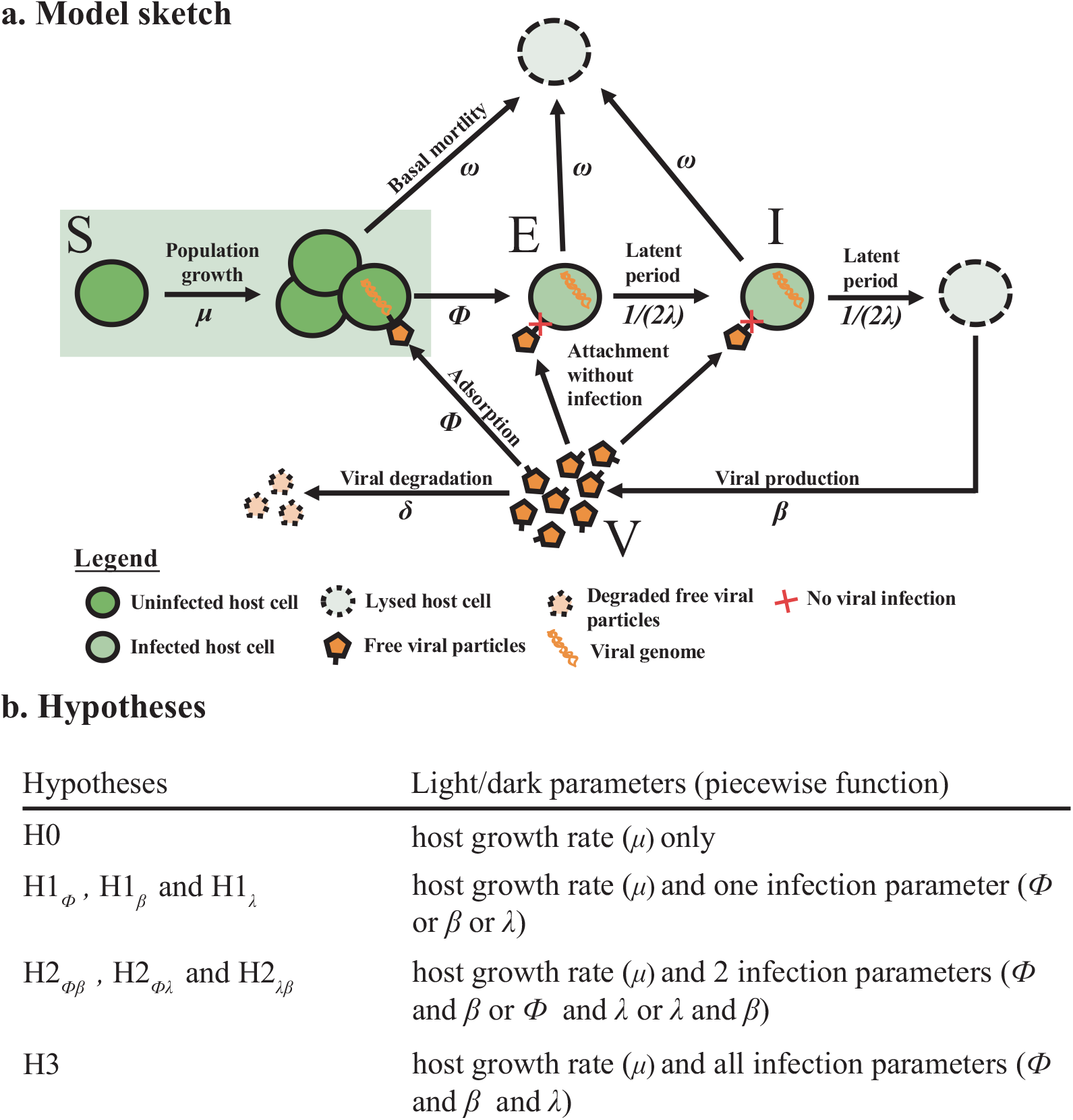
Description of the model. (a) Schematic representation of the model. The host population is divided into 3 classes: Susceptible to infection (S), Exposed (E) and Infected (I) by Virus (V). Black arrows are biological processes described by the mathematical parameters. (b) Definition of the hypotheses. Each hypothesis describes a possible relation between light and the infection parameters (orange) under different infection parameters sets (*θ_infection_*).

In this model, *μ* is the host growth rate (*h*^−1^), *K* is the host carrying capacity (cell mL^−1^), *ω* is the host basal mortality (*h*^−1^) not due to viral lysis, *ϕ* is the adsorption rate (mL *h*^−1^), *λ* is the latent period (h), *β* is the burst size (unitless), and *δ* is the viral decay rate (h^−1^) (see Supplementary Table II for more information on parameters). We assume that viruses can attach to all host cells (*S*, *E* and *I*) but only lead to state transitions when infecting *S* types, i.e., from susceptible to exposed.

We have already established that light modulates host growth (Figure 2). However, it is not evident if diel variation in host growth alone can explain changes in virus and host dynamics at population scales. Hence, we defined a series of nested model hypotheses that include alternative mechanisms for light-driven changes in viral life history traits (Figure 3b). The mechanisms are different in the number of viral life history traits that differ between light and dark. The number ranges from 0 (in the null hypothesis *H*0) to 3, where the adsorption rate, latent period, and burst size, each differ between light and dark. In practice, each model parameter that is light-driven takes on two values in the model, e.g., the burst size would have *β*_dark_ and *β*_light_. Although viruses are known to be degraded under UV light [19], our experiments were conducted under white light without UV radiation and viral decay rates were similar under light or dark conditions (Supplementary Figure 2 and Supplementary Table III). Henced, we fixed the value of decay rates.

We fit each of the nested, light-driven virus-host population models using MCMC to experimental measurements of *Prochlorococcus* strain MED4 infected by either cyanophage P-HM2 or P-SSP7 over a 4 day period (Figure 4). Parameter ranges in the MCMC procedure were constrained by prior estimates (Supplementary Figure 3 and Supplementary Table IV)[53]. We found the best fit model to be *H*2_*ϕλ*_ P-HM2 and *H*0 for P-SSP7. This suggests that P-HM2 (but not P-SSP7) has light-dependent life history traits that help provide explanatory power to the virus-host population dynamics. In both cases, viral abundances rapidly increase and then plateau. However, in both cases, the best-fit model significantly over-estimates the degree of viral-induced mortality in the host population, e.g., models predict that the final timepoint estimates of cell density are 2.5 and 6.1-times lower than measured for the P-HM2 and P-SSP7 cases, respectively. This result suggests that other features underlying interactions between cyanophage and *Prochlorococcus* have to be taken into account when scaling up to the population-level dynamics.

**FIG. 4:**
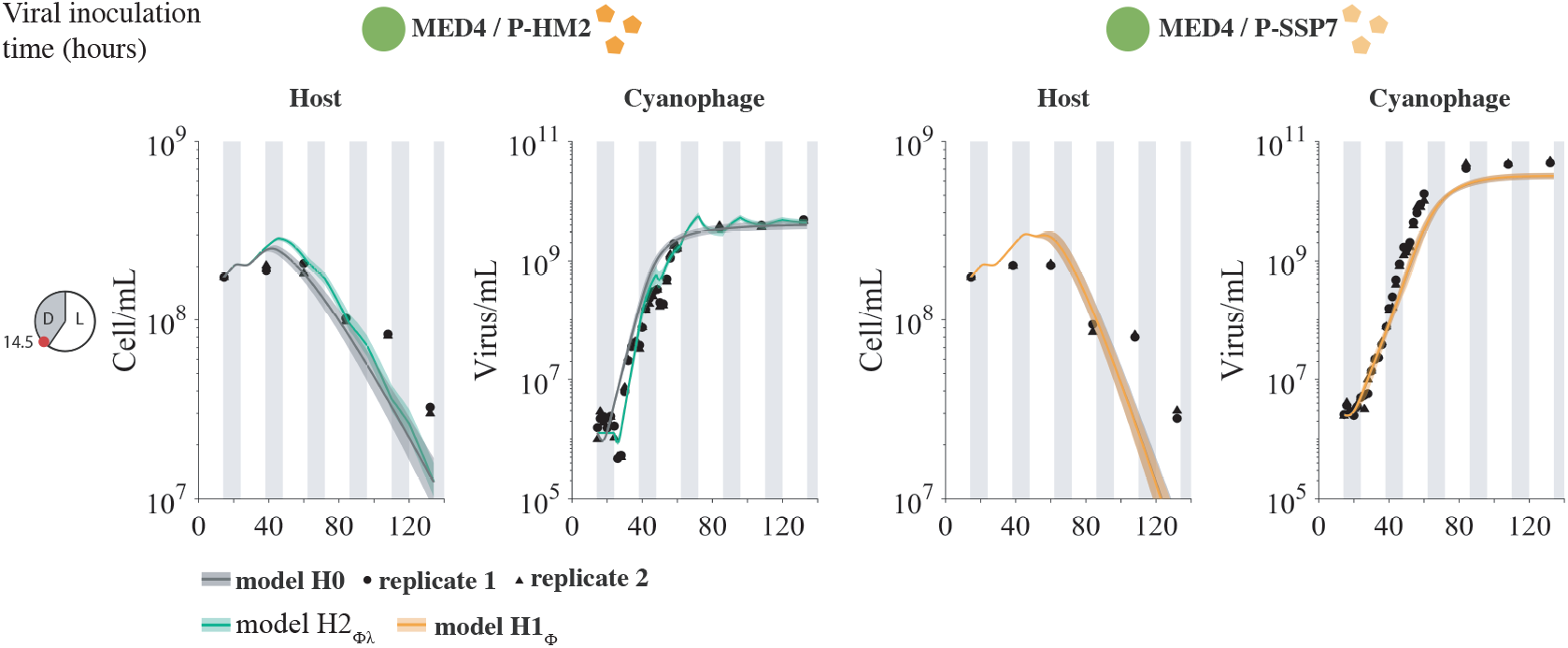
Light-driven models to host and virus population abundance data. Model fits under *H*0 (gray) and hypotheses *H*2_*ϕλ*_ (green) and *H*1_*ϕ*_ (orange) for an inoculation time of 14.5 hours after the beginning of the experiment. Phage P-HM2 infecting strain MED4 (left colones) and P-SSP7 infecting MED4 (right colones). Solid lines are the median of 5000 model simulations with shaded areas the 95% quantiles area. Data are represented by the black dots for two replicates. Vertical shaded gray lines represent dark conditions.

### Beyond light: incorporating lysis inhibition to explain virus and host population dynamics

The observation that host populations remain persistently above model expectations when viral abundances are high suggests a potential slowdown mechanism in viral-induced lysis. To account for this, we modified the initial model to account for an additional state transition, i.e., from *I* to *E*:

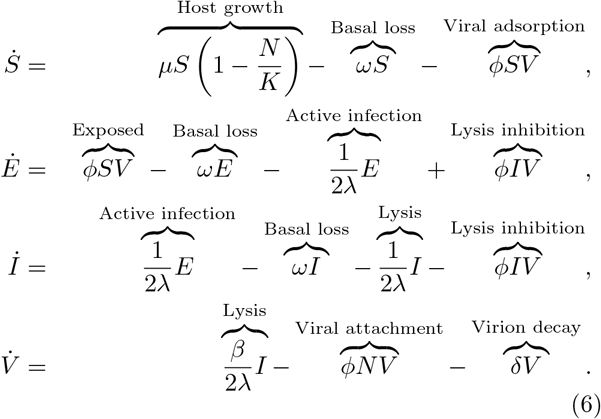

In this model, free virions switch the state of infection from *I* to *E*, thereby slowing down the expected time to lysis. This slow-down occurs in a fraction 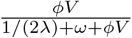 of cells in the *I* state; hence it increases with increasing virus density. For example, given the best-fit parameters for P-SSP7, this fraction changes from 1.28 10^−4^ when *V* = 10^6^ virions/ml to 1.26 10^−2^ when *V* = 10^8^ virions/ml; nearly a 100-fold difference. We denote Eq. (6) the lysis inhibition model.

We then fit the lysis inhibition model to an expanded set of experimental measurements of MED4 and infected by either cyanophage P-HM2 or P-SSP7. The measurements included addition of viruses at 14.5, 18, 24.5, 30 and 36 hrs after host addition (see Figure 5). The light-dependent hypothesis used in fitting are denoted as 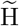 to distinguish them from the original hypotheses. Via an MCMC fitting procedure, we find that the models 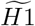 and 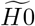 best fit the host and virus dynamics in the P-HM2 and P-SSP7 cases, respectively (Figure 5). Notably, best-fit model simulations are now able to reproduce both the viral saturation and the slowdown of the host population. A full list of AIC and BIC information is found in Supplementary Figure 4 and Supplementary Table V. Specifically, both P-HM2 and P-SSP7 can adsorb, replicate, and lyse cells in the light. However, models suggest that P-HM2 has markedly different light vs. dark infection life history traits, whereas there is not enough evidence to reject the null hypothesis in the case of P-SSP7.

**FIG. 5:**
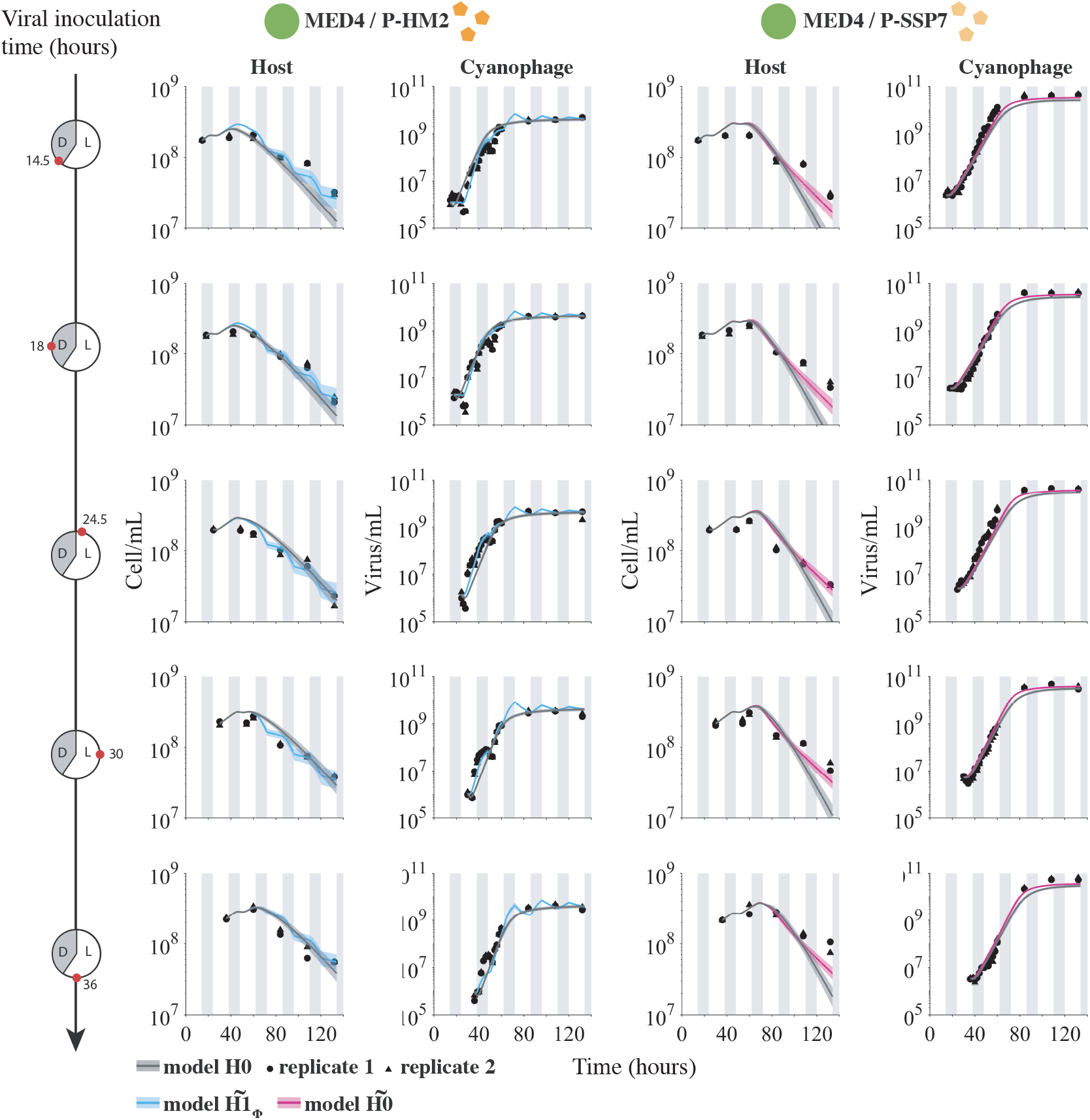
Viral dynamics under light-dark cycle for different viral inoculation times. Model fits under *H*0 (gray) and best hypotheses 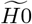 (pink) or 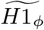 (blue) for different viral inoculation times. Phage P-HM2 infecting strain MED4 (left colones) and P-SSP7 infecting MED4 (right colones). Solid lines are the median of 5000 model simulations with shaded areas the 95% quantiles area. Data are represented by the black dots for two replicates. Vertical shaded gray lines represent dark conditions

We evaluated the quality of fits by assessing the predicted estimates of life history traits for the P-HM2 and P-SSP7 cases. Disparities in parameters under light or dark conditions obtained with our MCMC approach are consistent with earlier measurements of viral infections of MED4 give fixed light or dark conditions over a 10 hr period [37]. Specifically, model fits reveal that P-HM2 has a significantly lower adsorption rate in the dark compared to the light (Figure 6). Indeed, dark-adsorption is at the lower limits to the parameter constraint range of the MCMC procedure, suggesting that P-HM2 may have effectively zero adsorption in the dark. In contrast, model estimates cannot reject the hypothesis that adsorption was effectively constant for P-SSP7. The convergence of MCMC chains further support the robustness of the model-based inferences (see Supplementary Table IV, Supplementary Figures 5 and 6). Notably, other candidate models with intracellular mechanisms that delay lysis can also reproduce similar population-level features. For example, density-dependent changes in the development of the intracellular infection from the eclipse to the mature infection stage can also lead to a slowdown of lysis of the total host population (Supplementary Figure 7).

**FIG. 6:**
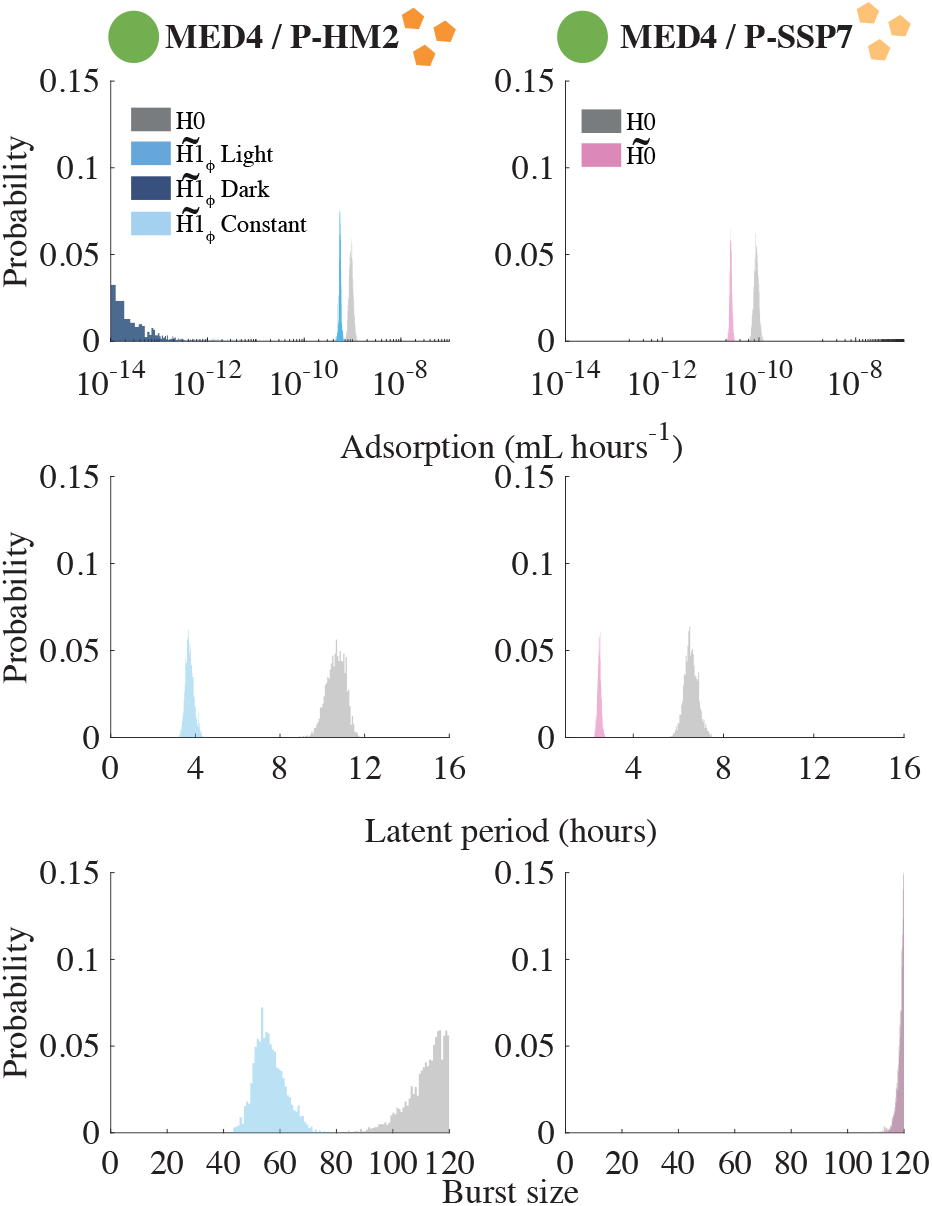
Infection parameter distributions: P-HM2 (left panels) and P-SSP7 (right panels) infecting strain MED4. Histogram distributions are calculated with 5000 parameter sets. Model parameters are represented in purple for dark and blue for light conditions and in gray when it is constant. Parameters are for the 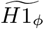 hypothesis for the pair P-HM2/MED4 (blue) and 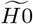 for P-SSP7/MED4 (pink) with comparison to the *H*0 hypothesis (gray).

## Discussion

We investigated the impact of light and dark conditions on the infection of *Prochlorococcus* by cyanophage, using a combination of experiments, nonlinear population models, and model-data integration. We found that light-dependent differences in viral adsorption to hosts help explain population-level changes in both virus and host abundances given growth in diurnal conditions. These light-dependent differences are strain-specific. Estimated adsorption rates vary markedly during the light vs. dark for P-HM2 but not for P-SSP7. This suggests that viruses, in addition to hosts, may have light-dependent differences in their life history traits at the cellular-scale that impact dynamics at population scales.

In our model-fitting procedure, we evaluated the possibility that light could affect adsorption, latent period, and burst sizes. Nonetheless, we only found evidence for a light-dependent variation in adsorption rate for the phage P-HM2. In contrast, P-SSP7 dynamics were better explained by light-driven host growth rate. Our results corroborate the observations of [37], adding mechanistic evidence of light-driven *Prochlorococcus* infection dynamics. The imputed failure of P-HM2 to adsorb to MED4 in the dark indicates that adsorption could be directly modulated by light, as suggested by [36]. Light-dependent variation in adsorption has also been reported in cyanophage infecting *Synechococcus* [36, 39] and Cocolithoviruses [48]. Notably, the work of [48] is most similar to our approach, in that it also used nonlinear models to identify light-driven variation in viral straits that impact virus-host population dynamics. For *Prochlorococcus*, there are multiple reasons why P-HM2 may have evolved such variation. First, exposure to UV is a critical factor degrading viral particles outside the host cell [19]. During the night, there is both less UV and (potentially) elevated predation rates of cyanobacteria by eukaryotic grazers ([54, 55]). Thereby, remaining outside the host cell during the night could effectively amount to a survival strategy by avoiding predation by grazers in the viral host [20]. Additional advantages may reflect context-specific advantages of partitioning adsorption differently depending on variation in host availability.

Despite our focus on light-driven traits, our approach revealed other mechanisms driving variation in host-virus population dynamics. The failure of a light-driven virus-host population model to recapitulate the persistence of host cells suggests additional feedback mechanisms that limit host mortality, even when virus densities are relatively high. Using a variant of the original model, we found evidence consistent with lysis inhibition at high viral densities (see [56]). Other possibilities that could explain this decrease in host decline include decreases in viral infectivity, an increase in the production of defective viral particles, or slowdowns in host physiology. Such slowdowns reflect the potential, reciprocal influence of processes at cellular and population scales. The relevance of such slowdowns will vary with environment. For example, in marine surface environments, cyanophage densities do not typically exceed 10^6^ ml^−1^, and so it remains unclear if the candidate feedback mechanism is an adaptive response to the high density of infected hosts, or arises incidentally given ecological conditions outside of typically encountered ranges. Further work is necessary to disentangle process from pattern.

In closing, we found that light-dependent viral life history traits can substantively change the dynamics of *Prochlorococcus* and cyanophage. This finding reinforces and extends the consequences of prior results showing that viral traits differ between light and dark, albeit in fixed conditions. In the marine environment, adaptation to light has been shown to drive differences amongst *Prochlorococcus* physiology as well as evolutionary adaptation between light-associated ecotypes. Our study suggests that exploring variation in viral-associated light-dependent life history traits may also reveal ways in which viruses partition their environment, both in terms of host specificity and via differential infection of hosts over light-dark cycles.

## Material and Methods

### Experimental design and data attribution

Experimental data analyzed here is from two sources: Liu *et al.* 2017 [51] and Liu *et al.* 2019 [37], and new data collected to link infection-level dynamics with population-level dynamics. Specifically, the host growth data in Figure 2 was previously reported in Liu *et al.* 2017 [51]. The infection data in Figure 1C and 1D as well as the host and phage abundances before 60 hrs for Figures 4 and 5 are reported in Liu *et al.* 2019 [37]. The host and phage abundance data after 60 hrs in Figures 4 and 5 as well as the viral decay decay reported in Figure S7 is newly reported here. Details of the experimental procedures are described in the following sections with full quotations used denoting when methods are equivalent to those reported in [37]. We include full methods descriptions for completeness.

#### Culture conditions

As in [37]: “Axenic *Prochlorococcus* strains were grown in Port Shelter (Hong Kong) seawater-based Pro99 medium [57]. Batch cultures were incubated at 23°C in continuous light (25 *μ*mol quanta m^−2^ s^−1^) or a 14h light:10h dark cycle (35 *μ*mol quanta m^−2^ s^−1^ in the light period). Cultures were acclimated in the same condition for at least three months before they were used for the experiments.”

#### Infection of synchronized Prochlorococcus cells under light-dark cycles

As in [37]: “*Prochlorococcus* cells were acclimated under light-dark cycles for at least three months and were synchronized, as determined by flow cytometry. Mid-log cells were infected at different times of a light-dark cycle at a phage/host ratio of 0.02. Times of infection were 0.5, 6, 12 hours. Each experiment was replicated at least two times.”

#### Cyanophage decay rates under light or dark

To measure the decay rates, fresh lysates of cyanophages P-HM2 and P-SSP7 were prepared by adding 300 *μ*L virus stocks into 30 mL mid-long *Prochlorococcus* MED4 culture. After the infected culture became clear, cell debris was removed using a 0.2 *μ*m polycarbonate filter and the supernatant containing phage particles was stored at 4°C in the dark. During the viral decay experiment, aliquots of viral lysates were incubated at 23°C at a light intensity of 27 *μ*mol photons m^−2^ s^−1^, and aliquots were incubated at the same temperature in the dark [58]. Samples were taken from each tube every two days over 10 days. To measure the loss of viral infectivity, the number of plaque forming units was measured [59]. Briefly, 500 *μ*L serially diluted viral lysate was added to 2 ml *Prochlorococcus* MED4 (grown to mid-log phase in Pro99) in glass tubes and incubated at room temperature for 15 min to allow phage adsorption. Incubated cultures were then combined with UltraPureTM low-melting-point agarose (Invitrogen) at a final concentration of 0.5%. The EZ55 Alteromonas helper bacterium [60] was added to every plate. Plaques began to appear seven days later on plates that were incubated at 23°C at the light intensity of 19 *μ*amol photons m^−2^ s^−1^. Each sample was measured with three technical replicates.

#### Flow cytometry and cell cycle analysis

As in [37]: *“Prochlorococcus* cells were preserved by mixing 100 *μ*L culture with 2 *μ*L 50% glutaraldehyde to a final concentration of 1% and were stored at −80°C. Cells were enumerated by a flow cytometer (BD FACSCalibur) with the CellQuestPro software. We followed a published protocol to determine the percentage of cells in each cell cycle stage [21]. Briefly, *Prochlorococcus* cells were stained with the DNA stain SYBR Green (Invitrogen) and flow cytometry data were analyzed with the ModfitLT software.”

#### Quantification of cyanophage

As in [37]: “Total phage particles were collected on a 0.02 *μ*m Whatman Anodisc filter, stained with SYBR gold (Molecular Probes), and counted under an epiflourescence microscope [61, 62]. At least five discrete fields on a filter were photographed using the SPOT Advanced Imaging software and fluorescent dots representing phage particles were counted manually.”

“During infection, extracellular phage DNA was quantified using a quantitative polymerase chain reaction (qPCR) method [46]. Briefly, infected *Prochlorococcus* cultures were filtered through 0.2 *μ*m polycarbonate filters in a 96-well filter plate (Pall). Filtrates containing extracellular phage particles were diluted 100-fold in dH2O and were then used as templates for qPCR reactions in a 384-well plate. A qPCR reaction contained 4.6 *μ*L template, 0.2 *μ*L forward primer (10 *μ*M), 0.2 *μ*L reverse primer (10 *μ*M), and 5 *μ*L iTaq Universal SYBR Green Supermix. The LightCycler 480 Real-Time PCR System (Roche Diagnostics) was used for thermal cycling, which consisted of an initial activation step of 5 min at 95° C, 45 amplification cycles of 10 s at 95° C and 60 s at 60°C, and a melting curve analysis at the end. The number of cyanophage in each well was quantified using a standard curve generated from phage particles that were enumerated by epifloures-cence microscopy. qPCR Phage DNA copies measured provides a ~ 1:1 relationship with phage particle counts [53].”

### Model simulations

Model analyses were performed with Matlab version 9.2.0 (Natick, Massachusetts: The MathWorks Inc., 2017). Infection dynamics were simulating using Matlab ODE solver *ode*45 [63](Copyright 1984-2011 The MathWorks, Inc.) which uses a higher-order Runge-Kutta method [64].

### Estimation of the best parameter sets

#### General procedure

The parameters estimation aimed to estimate the best model parameters set *θ* associated to the fits describing the measurements of hosts and viruses. To estimate the best parameters set, we decomposed the procedure into two steps: First, we estimated host growth parameters *θ_host_* for the model without viruses (Eq. 1) and, in a second time, we used *θ_host_* in the model with viruses (Eq. 5 and 6) and estimated the infection parameters *θ_infection_* for each hypotheses. The general procedure aimed to minimize an objective function that calculates the model fits error with experimental data for a given parameter set.

#### The objective function

The objective function calculated error between the model fits and the measurements as following (Eq. 7):

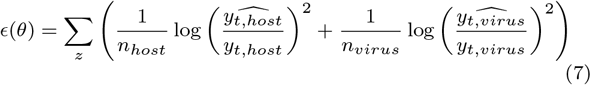

where *ϵ* is the total calculated error for the given parameter set to be estimated *θ*, over the *z* experiments. We decomposed the error into host and virus with *y_t,host_* being the host data at time 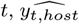 being the host model estimate at time *t* such that 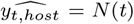 where *N*(*t*) is the sum of the susceptible, exposed and infected host cell estimations. Similar notations were used for the virus error. Then the total error result was the sum of the host and virus errors for the whole set of experiments.

#### Algorithms

We first sampled the parameters space with a Latin Hypercube Sampling (LHS) [65] for 20.000 parameter sets for each hypotheses and models. We then calculated the initial errors by running the model with these parameter sets and calculated the objective function. In a second time, we implemented a Markov Chain Monte Carlo (MCMC) procedure for the 10 best initial parameters in order to converge to the global minimum after two burn-in periods (running periods that allow the convergence of the chains). The final chains represented the distributions of the parameters for 5000 sets that minimize the objective function. We used the MCMC toolbox for Matlab implemented the DRAM algorithm [66].

#### Estimation of the host growth parameters θ_host_

For the host parameters set *θ_host_* = (*α, L_opt_, μ_max_, k_d_, ωandK*) we used the procedure described previously to estimate the parameters describing the growth of *Prochlorococcus* strains MED4. Parameters of the growth-irradiance curves (*α, L_opt_* and *μ_max_* – Eq. 4), were constrained by the data from Moore and Chisholm 1999 [52] whereas the carrying capacity *K* was fixed and considered as a constant (K = 3.10^9^ cell mL^−1^ - Unpublished data).

#### Estimation of the infection parameters θ_infection_

To estimate the parameter set *θ_infection_* = (*ϕ, λ, β*), we fixed the host growth parameters estimated previously and estimated the parameters relative to the hypotheses *H*0 to *H*7 described in Figure 2b. Depending on the hypothesis, the estimated parameter could be constant during the experiments (no relation with light or dark condition) or piecewise function (light and dark values). The estimated parameters were the adsorption rate *ϕ*, the latent period *λ* and the burst size *β*. Viral decay rates were estimated experimentally as the slope of log(viral concentration) regression under light or dark conditions and fixed (see Supplementary Figure 2 and Table III).

### Quantifying the best model hypothesis

To quantify the best model under hypothesis *H*0 to *H*7 we computed an Akaike Information Criterion (AIC) and a Bayesian Information Criterion (BIC) [67], for virus and host (when data were available) according to the following equations (Eq. 8, 9):

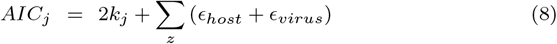

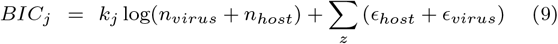

with 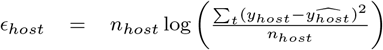 and 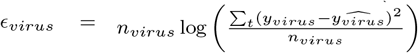. These criteria are computed depending on the hypothesis *j* with the number of parameters to be estimated *k_j_* (3 parameters for *H*0, 4 for hypotheses *H*1, *H*2 and *H*3, 5 for hypotheses *H*4, *H*5 and *H*6 and 6 for hypothesis *H*7 – Figure 2b), *n_host_* and *n_virus_* being the total number of data points for host and virus respectively, *z* being the treatment, *y_host_* and *yvirus* being the data points for the hypothesis *j* and the treatment *z* at time point *t* for host and virus respectively, 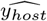 and 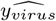 being the model estimation for the hypothesis *j* and the treatment *z* at time point *t* for host and virus respectively.

### Data availability statement

All data is available for use and re-use. The full dataset and code is available at `DOI:10.5281/zenodo.3308790`. As noted in the Methods, the analysis here includes data from both published and unpublished sources. Data for Figure 1c and 1d are from [37]. Data for Figure 2a are from [51]. Data for Figure 4 and 5 until 60 hours are reused from [37] and new for measurements after 60 hours. Data for Supplementary Figure 2 are original.

## Acknowledgments

This study was supported by grants to Qinglu Zeng from the National Natural Science Foundation of China (Project number 91851112) and the Research Grants Council of the Hong Kong Special Administrative Region, China (Project number 16144416) and by a grant to Joshua Weitz from the Simons Foundation (SCOPE Award ID 329108). We thank Akram Salam for feedback on the manuscript.

## Author contribution

DD, QZ, and JSW designed the study. RL, YC, and FZ performed experiments, designed by QZ. DD carried out the modelling analysis with contributions of JSW. DD and YC performed candidate model analysis. AC contributed to code development. DD and JSW wrote the manuscript with contributions from QZ.

## SUPPLEMENTARY MATERIAL

### Supplementary Text - Model Analysis

We tested different hypotheses and built candidate models based on the initial model that can explain the lysis slowdown. Here we listed the candidate models that reproduced a lysis slowdown and represent their dynamics in the Supplementary Figure 7.

#### Candidate model 1: different eclipse and lysis periods

We used the initial model (5) and subdivided latent period into two periods: the eclipse period *λ_e_* and the lysis period *λ_l_* such that *λ* = *λ_e_* + *λ*;. The dynamics of the candidate model 1 are represented with cyan solid lines in the Supplementary Figure 7.

#### Candidate model 2: lysis decrease with viral concentration

The candidate model 2 is similar to the initial model (eq. 5) but the latent period is now a function of the viral concentration as follows:

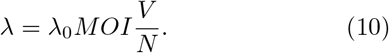

where *λ*_0_ is the basal latent period and *MOI* is the MOI rate of the treatment (*MOI* = 0.1). The dynamics of the candidate model 2 are represented with green solid lines in the Supplementary Figure 7.

#### Candidate model 3: host cells mutation lead to resistance to infection

The candidate model 3 is based on the initial model (eq. 5) but allows the host cell *S* to mutate at a rate *r* and become resistant to infection. The resistant host cells *R* grow at a lower rate than *S*, given a presumed fitness cost to resistance:

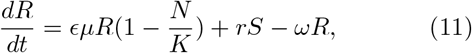

where *ϵ* is the cost of resistance (*ϵ* = 0.5), *N* is the total host population such that *N* = *S* + *E* + *I* + *R* and *r* is the mutation rate from the class S to R. We used the mutation rate measured by [68] for *Prochlorococcus* strain MED4 and consider that cell divide ones per day (*r* = 6.08 10^−6^ per division). The dynamics of the candidate model 3 are represented with pink solid lines in the Supplementary Figure 7.

#### Candidate model 4: unsuccessful adsorption

The candidate model 4 is based on the initial model (eq. 5) but we assume that adsorption does not necessarily lead to infection. Instead, viruses *V* can adsorb at a rate *ϕ* to susceptible cells *S* which leads to a new infection in an *ϵ* fraction of cases. The dynamics of candidate model 4 are represented with red solid lines in the Supplementary Figure 7.

**SI. TAB. I:** Host growth parameters means and standard deviations.

| Parameters | MED4 |
| --- | --- |
| $\alpha$ | $0.0011 \pm 3.10 \cdot 10^{-4}$ |
| $L_{opt}$ | $44.78 \pm 7.98$ |
| $\mu_{max}$ | $0.035 \pm 0.0021$ |
| $K_L$ | $151.59 \pm 17.51$ |
| $\omega$ | $0.0032 \pm 0.0012$ |

**SI. TAB. II:** Description of model parameters

| Parameters | Meaning | unit |
| --- | --- | --- |
| $\mu$ | Host growth rate | $h^{-1}$ |
| $K$ | Host carrying capacity | Cell $mL^{-1}$ |
| $\omega$ | Host mortality | $h^{-1}$ |
| $\phi$ | Viral adsorption | $ml \ h^{-1}$ |
| $\lambda$ | Viral lysis | $h^{-1}$ |
| $\beta$ | Burst size | no dimension |
| $\delta$ | Viral decay | $h^{-1}$ |

**SI. TAB. III:** Experimental estimation of viral decay values using plaque assay.

| Phage | $\delta$ light ($d^{-1}$) | $\delta$ dark ($d^{-1}$) |
| --- | --- | --- |
| P-HM2 | 0.022 | 0.0289 |
| P-SSP7 | 0.0176 | 0.021 |

**SI. TAB. IV:**
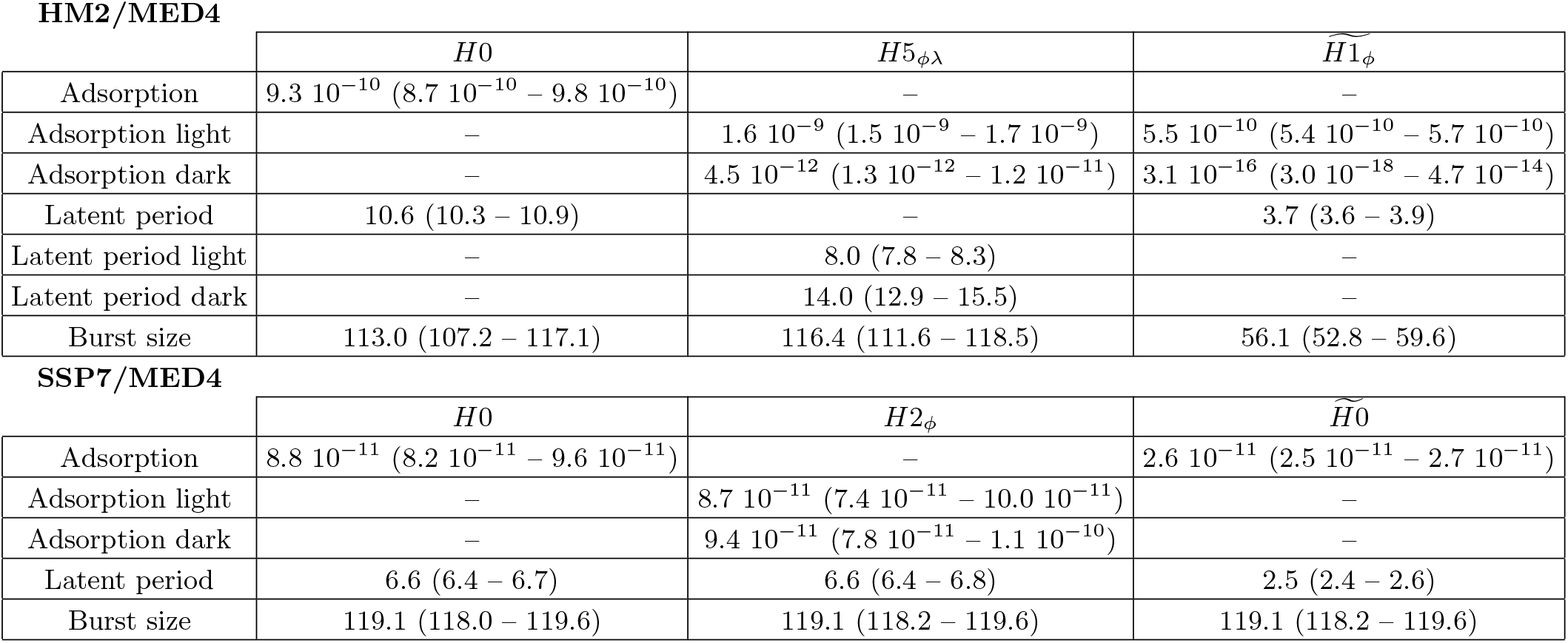
Statistics: median and quantiles (25% – 75%) of the parameter distributions for best initial and lysis inhibition model hypotheses for virus P-HM2 and P-SSP7 infecting *Prochlorococcus* strain MED4, calculated with 5000 parameter sets.

**SI. FIG. 1:**
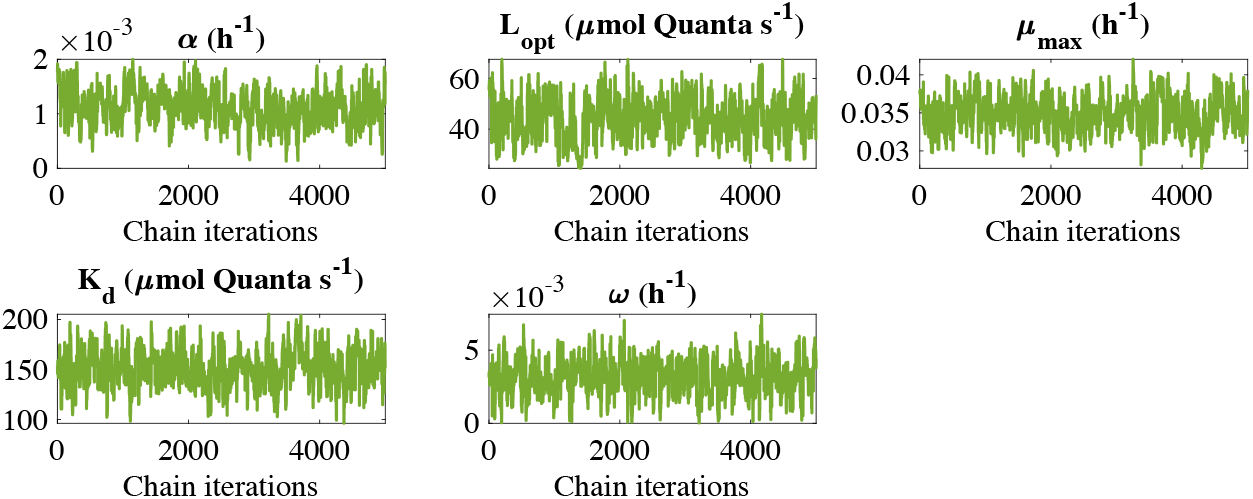
Chain convergences for the host growth parameters for *Prochlorococcus* strain MED4.

**SI. FIG. 2:**
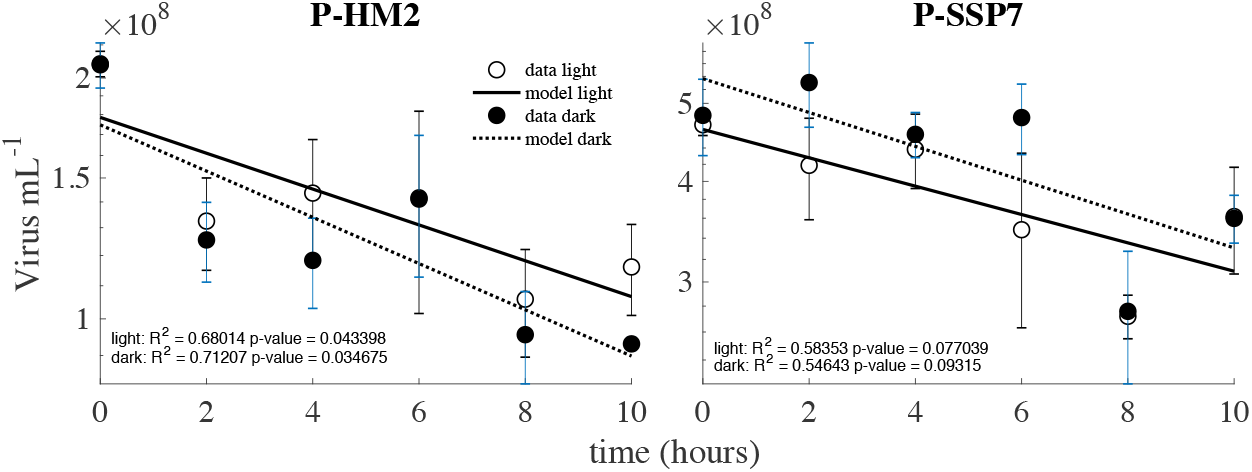
Experimental estimation of viral decay in light and dark condition for P-HM2 and P-SSP7.

**SI. TAB. V:**
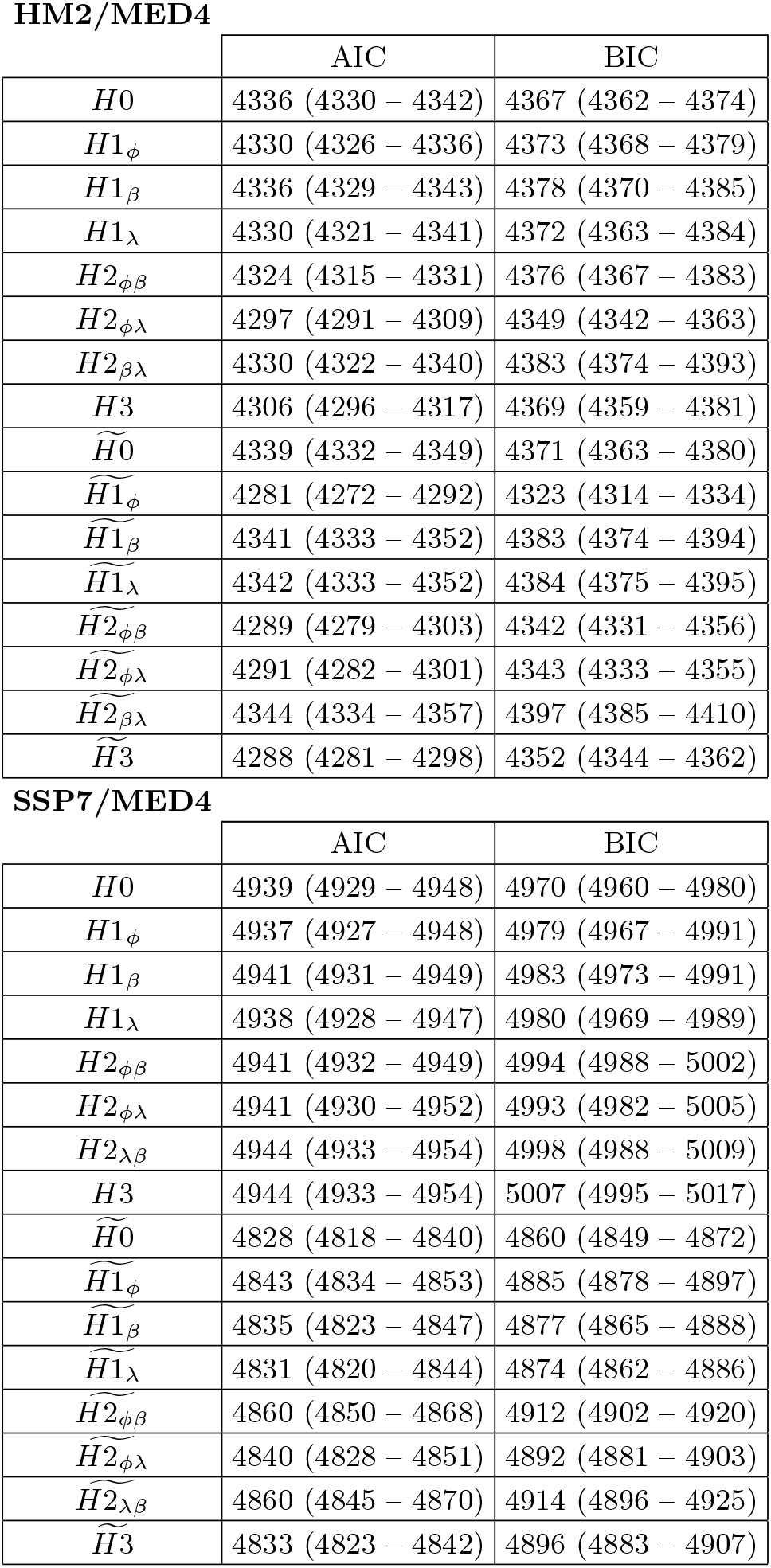
Akaike Information Criteria (AIC) and Bayesian Information Criteria (BIC) median and quantiles (25% – 75%) values for 5000 parameters sets. Minimum value are corresponded to the best model.

**SI. FIG. 3:**
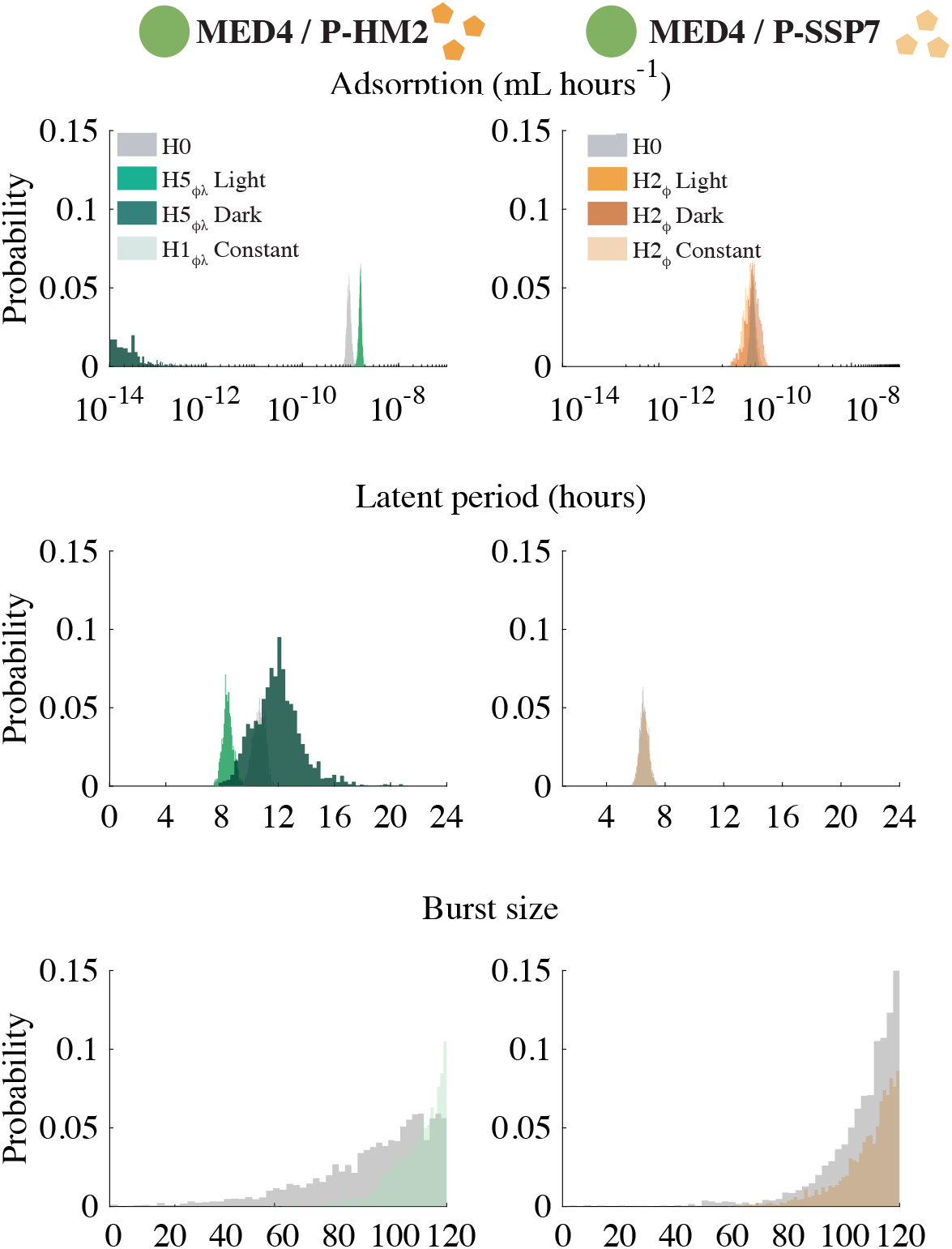
Best initial model infection parameter distribution: P-HM2 (left panels) and P-SSP7 (right panels) infecting strain MED4. Histogram distributions are calculated with 5000 parameters sets.

**SI. FIG. 4:**
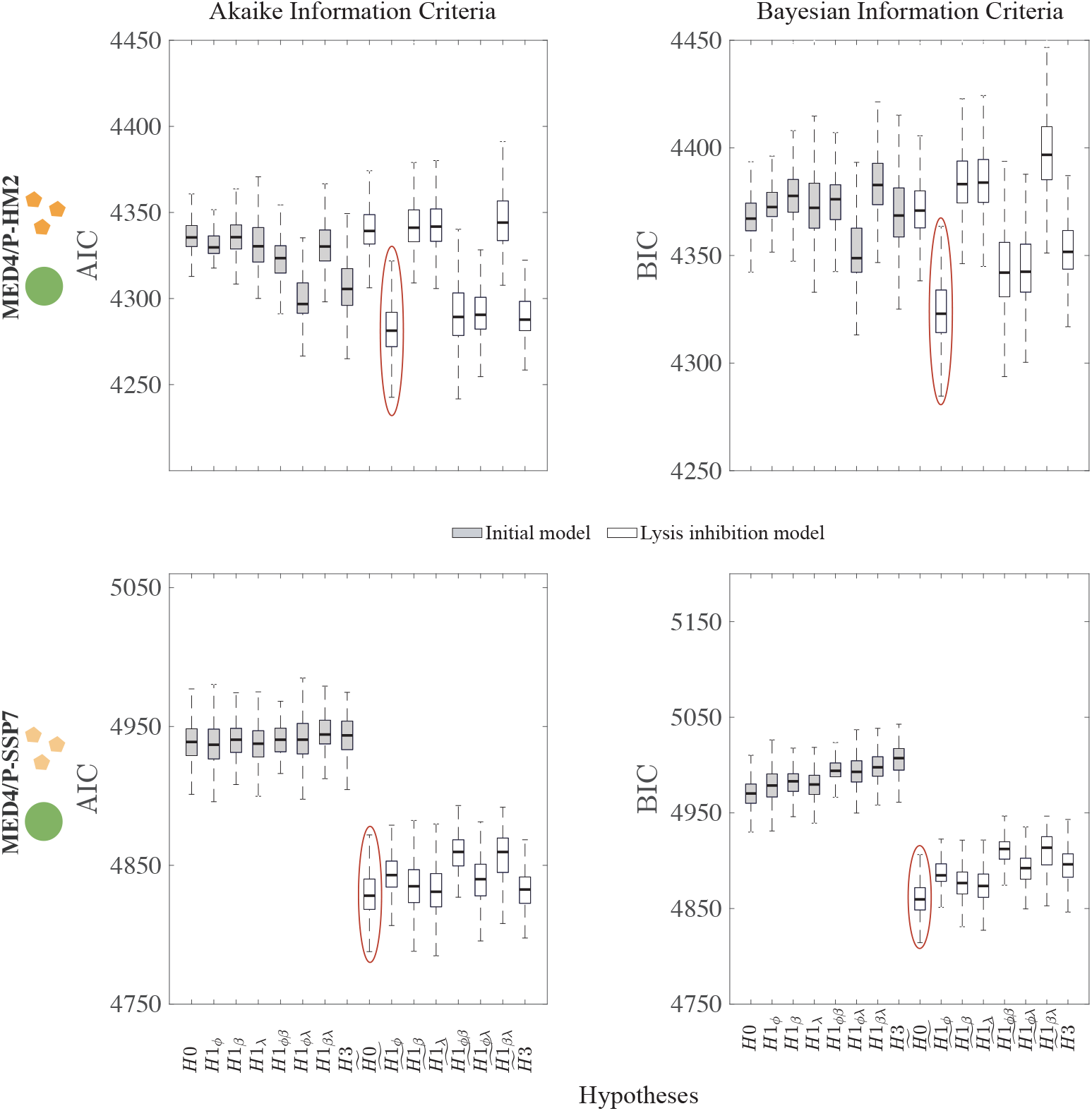
Model comparison criterion. Akaike Information Criteria (AIC - left panels) and Bayesian Information Criteria (BIC - right panels) calculated for the 2 Host-virus pairs used in the study. Box plots are calculated over 5000 parameters sets for each hypothesis and model. Minimum values of AIC and BIC considering the best hypotheses. Red circles indicate the best hypotheses for each host-virus pairs.

**SI. FIG. 5:**
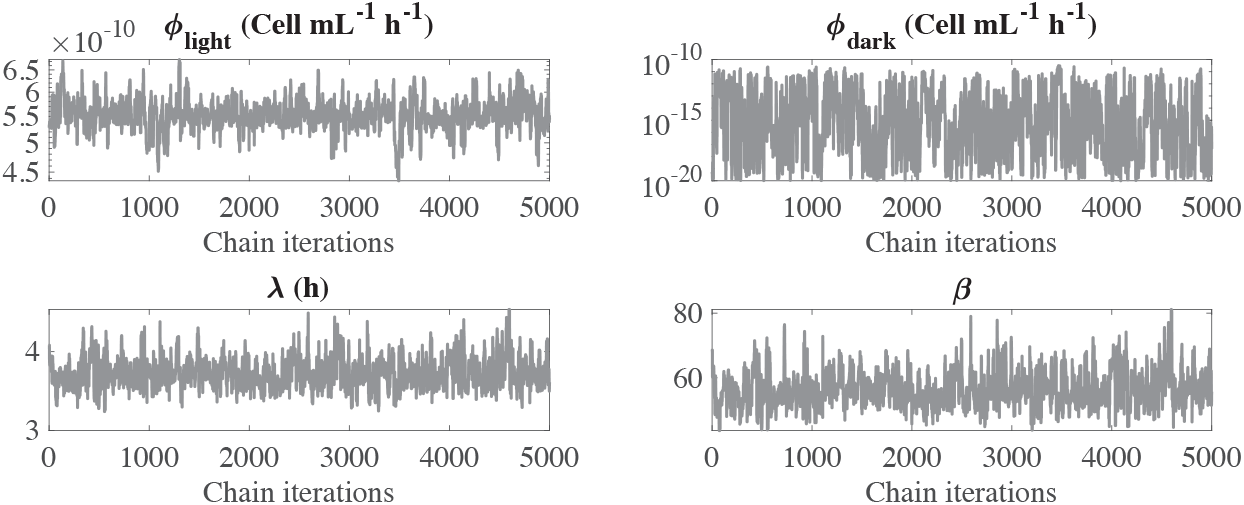
Chains convergences for infection parameters of P-HM2 infecting strain MED4 for the best hypothesis 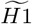.

**SI. FIG. 6:**
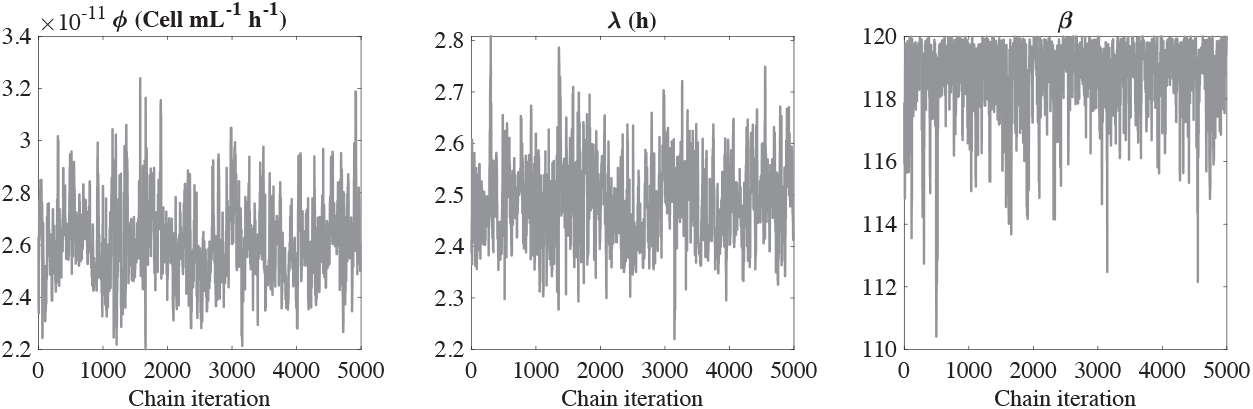
Chains convergences for infection parameters of P-SSP7 infecting strain MED4 for best hypothesis 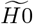.

**SI. FIG. 7:**
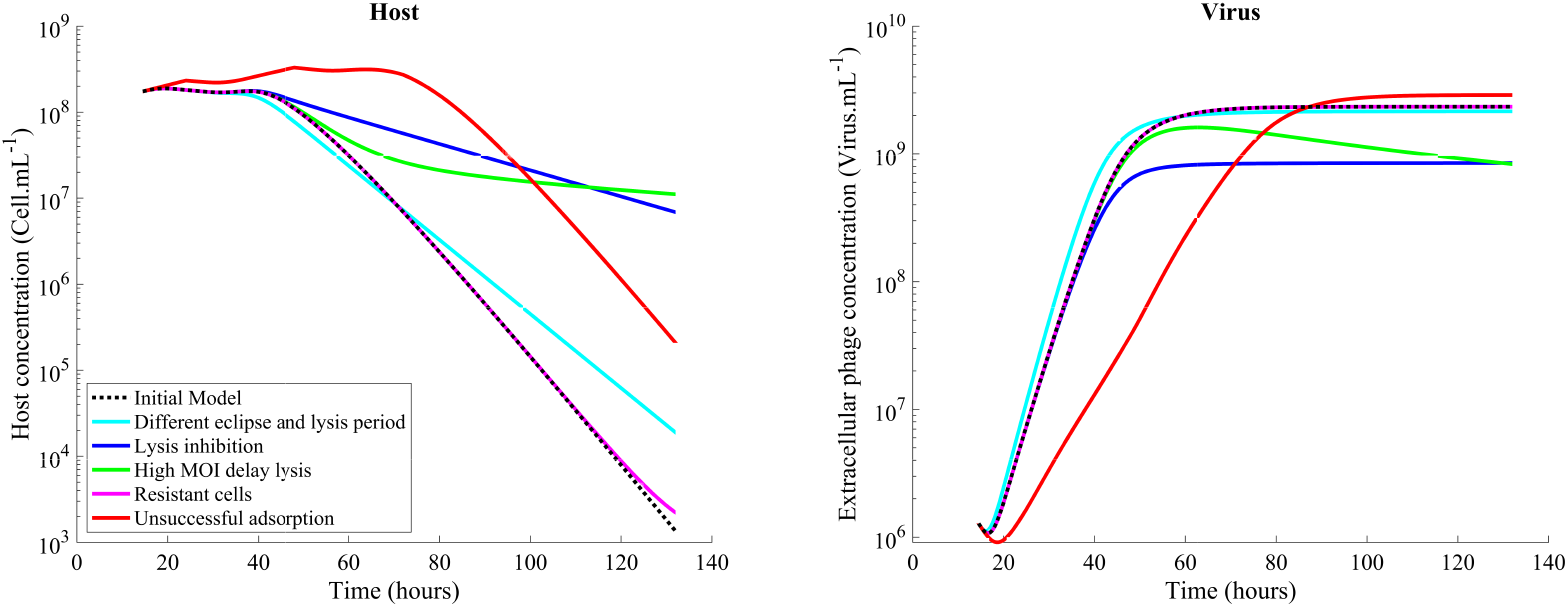
In silico experiments with candidate models that reproduce a slowdown of the population lysis. The initial model is represented by dashed black line, and the candidate models are in solid colored lines.

